# Mitochondrial respiration is essential for photosynthesis-dependent ATP supply of the plant cytosol

**DOI:** 10.1101/2024.01.09.574809

**Authors:** Antoni M. Vera-Vives, Piero Novel, Ke Zheng, Shun-ling Tan, Markus Schwarzländer, Alessandro Alboresi, Tomas Morosinotto

## Abstract

Plants rely on solar energy to synthesize ATP and NADPH for photosynthetic carbon fixation. Since a substantial proportion of cellular ATP is consumed in the cytosol, photosynthesis-derived ATP needs to be supplied there. While the triose phosphate shuttle and mitochondrial respiration can both deliver ATP to the cytosol, the significance of the different mechanisms *in vivo* has been difficult to assess. Although mitochondrial respiration is essential in plants, whether this is due to heterotrophic bottlenecks during plant development or rather a need for respiration in photosynthetically active cells, has not been resolved. In this study, we examined *in vivo* changes of cytosolic ATP concentration in response to light, employing a biosensing strategy in the moss *Physcomitrium patens*. Our measurements revealed increased cytosolic ATP concentration caused by photosynthetic activity. Moss tissue depleted of respiratory complex I showed decreased cytosolic ATP accumulation, highlighting a critical role of mitochondrial respiration in light-dependent ATP supply of the cytosol. Consistently, targeting mitochondrial ATP production directly, through the construction of mutants deficient in mitochondrial ATPase (complex V), led to drastic growth reduction, despite only minor alterations in photosynthetic electron transport activity. Since *P. patens* is photoautotrophic throughout its development, we conclude that heterotrophic bottlenecks cannot account for the indispensable role of mitochondrial respiration in plants. Instead, our results offer compelling evidence that mitochondrial respiration is essential for ATP provision to the cytosol in actively photosynthesizing cells. Mitochondrial respiration provides metabolic integration, ensuring a reliable supply of cytosolic ATP essential for supporting plant growth and development.

## Introduction

Photosynthetic organisms are the main primary producers on our planet, fixing approximately 110 Gt Carbon per year and providing the chemical energy supporting most lifeforms (Ringsmuth et al., 2016). Sunlight powers the photosynthetic electron flow catalysed by photosystem (PS) I and II, cytochrome *b_6_f* and the ATP synthase that together mediate the synthesis of NADPH and ATP to drive cellular metabolism. All eukaryotic photosynthetic organisms require mitochondrial respiration in addition to photosynthesis. Respiration allows for the transfer of electrons from metabolic intermediates to oxygen through the activity of the respiratory complexes localized in the inner mitochondrial membrane, complex I, II, III and IV, as well as alternative electron transport proteins, such as NAD(P)H dehydrogenases and alternative oxidases. Electron transport via complexes I, III and IV is coupled to proton transport and the generation of an electrochemical gradient across the inner mitochondrial membrane that drives the synthesis of ATP through the F_1_F_O_ ATP synthase, also called complex V. The NADH:ubiquinone oxidoreductase complex (complex I, CI) is under most conditions a major site of electron entry into the mitochondrial electron transport chain (mETC) and it can provide up to 40 % of the protons exploited for mitochondrial ATP synthesis (Watt et al., 2010; Braun et al., 2014), even though this varies widely depending on metabolic status and the contribution of alternative electron transport components.

Respiration in plants is essential for survival. Functional knockouts of respiratory complexes II, III, IV and V lead to lethality (León et al., 2007; Robison et al., 2009; Radin et al., 2015; Kolli et al., 2020). One evident explanation that applies for many plants is the fact that mitochondria cover cellular energy demand during the night and in non-photosynthetic tissues, such as roots. Importantly, seed plants typically go through developmental stages where photosynthesis is not active, such as embryogenesis and seed germination. While the need for energy supply in the absence of photosynthetic activity provides a potential explanation for why respiration is essential in plants, an increasing body of evidence suggests a relevant role of respiration also for sustaining photosynthesis (Noguchi and Yoshida, 2008; Joliot and Joliot, 2008; Bailleul et al., 2015; Burlacot and Peltier, 2023). A tight functional link between chloroplast and mitochondrial energy metabolism has been suggested (Dutilleul et al., 2003; Cardol et al., 2003; Schönfeld et al., 2004; Mellon et al., 2021). In the diatom *Phaeodactylum tricornutum* it was demonstrated that metabolite exchange between chloroplast and mitochondria is essential for carbon fixation (Bailleul et al., 2015). Moreover, excess reducing power produced via photosynthesis can be routed to mitochondrial respiration, preventing over-reduction of electron transporters and generation of reactive oxygen species (ROS) in the plastid (Noguchi and Yoshida, 2008; Zhang et al., 2012).

While only knock-down plants have been isolated and studied for complexes II, III, IV and V so far, plants completely lacking mitochondrial Complex I activity have been described in *Arabidopsis thaliana* as well as in *Nicotiana sylvestris* and the moss *Physcomitrium patens* where they showed a severe growth phenotype and, in the formers, alterations in germination, fertilization, and pollen development (Gutierres et al., 1997; Fromm et al., 2016; Mellon et al., 2021). Interestingly, European mistletoe *Viscum album* was shown to be able to live without a functional CI and most of the relevant corresponding genes have been lost (Maclean et al., 2018; Senkler et al., 2018; Schröder et al., 2022). While this is an extreme example of evolutionary rearrangement of central energy metabolism, probably linked to the specific lifestyle as an obligate semi-parasite (although currently unique to mistletoes and not found in other parasites), also *Viscum* shows respiratory activity, likely based on alternative NADH dehydrogenases.

In contrast to plants, a range of respiratory mutants depleted in all respiratory complexes has been isolated in the green alga *Chlamydomonas reinhardtii*. Those mutants generally show strong phenotypes under heterotrophic conditions but grow similarly to WT under photoautotrophic conditions (Salinas et al., 2014; Larosa et al., 2018). Those observations may be taken as evidence that mitochondrial respiration is only strictly required to support heterotrophic phases and is not essential when photosynthesis is active. Alternatively, there may be a fundamental difference in the role that respiration plays in plants and algae that may have evolved to making plant, but not algal, photosynthesis strictly dependent on mitochondrial respiration (Mellon et al., 2021).

In photosynthetically active cells, both the chloroplasts and the mitochondria are able to synthesize ATP. ATP is, however, also essential in other subcellular compartments. In the cytosol major ATP consumption occurs to support protein synthesis, membrane transport or primary metabolic pathways. It was proposed that when chloroplasts actively reduce CO_2_, cytosolic ATP originates mainly from the mitochondria as exported directly via the ATP-ADP-carriers (AACs). In contrast, ATP export from chloroplast was suggested to be enhanced under stress conditions, when CO_2_ fixation is limited (Gardeström and Igamberdiev, 2016). Even in that case ATP export would be indirect via the triose-phosphate shuttle, since there is currently no evidence for any direct and substantial ATP export from the stroma to the cytosol. These conclusions derive from observations made from experiments using biochemical fractionation and inhibitors and were more recently supported also by modelling approaches (Shameer et al., 2019). Yet, information from intact systems, that enable the exploration of dynamic transitions and responses to variable environmental conditions are largely missing, which represents a remarkable shortcoming in our current understanding of energy metabolism considering the central role of ATP for (plant) life.

In this work we set out to address the question why mitochondrial respiration is essential in plants. Specifically, we aimed at distinguishing between the two hypotheses that either (i) mitochondrial respiration is essential due to the need to sustain heterotrophic cells and heterotrophic phases of development where photosynthesis is absent, or (ii) mitochondrial respiration is critical for cellular ATP supply in the presence of photosynthesis. We chose the moss *P. patens* as a model that is able to complete its life cycle entirely without any heterotrophic developmental stage, enabling isolation of mutants that are not viable in other plants. Further, we addressed the methodological limitation of insufficient subcellular resolution in ATP measurements by exploiting a genetically encoded FRET sensor (ATeam1.03-nD/nA) that allows monitoring of rapid ATP changes as recently demonstrated in Arabidopsis (De Col et al., 2017; Voon et al., 2018; Lim et al., 2022). To measure the impact of photosynthetic activity on ATP *in situ* we further adapted an on-stage illumination approach (Elsässer et al., 2020). Exploiting cytosol-specific ATP monitoring in *P. patens* enabled us to explore the role of the mitochondria in cytosolic ATP supply. Genetic inactivation of respiratory complexes I and V allowed us to gather compelling evidence that mitochondrial respiration is essential for cytosolic ATP supply also while photosynthesis is active.

## Results

### Generation of P. patens plants accumulating ATP probe ATeam in the cytosol

Stable *P. patens* wild-type (WT) lines constitutively accumulating ATeam1.03-nD/nA in the cytosol, referred to as WT-ATeam, were generated by protoplast transformation. After transformation, lines stably resistant to zeocin were screened through the fluorescence of the mVenus channel to select the ones with detectable accumulation of the probe. At least three independent lines were selected among the ones showing mVenus fluorescence signal (Figure S1A, B). The ATeam-expressing lines did not show any visible growth defect and showed normal development (Figure S1A, B) with the ability to grow leaflets, rhizoids and sporophytes and produce viable spores as well as the parental line, suggesting that the probe accumulation did not cause major alterations in metabolism and development.

The localization of the mVenus fluorescent signal in the lines was verified by confocal microscopy, with chlorophyll fluorescence marking chloroplast localization. The mVenus signal showed no overlap with chlorophyll fluorescence and was localized in large areas inside the cell, consistent with a cytosolic expression, confirming the expected localization (Figure 1A).

**Figure 1.**
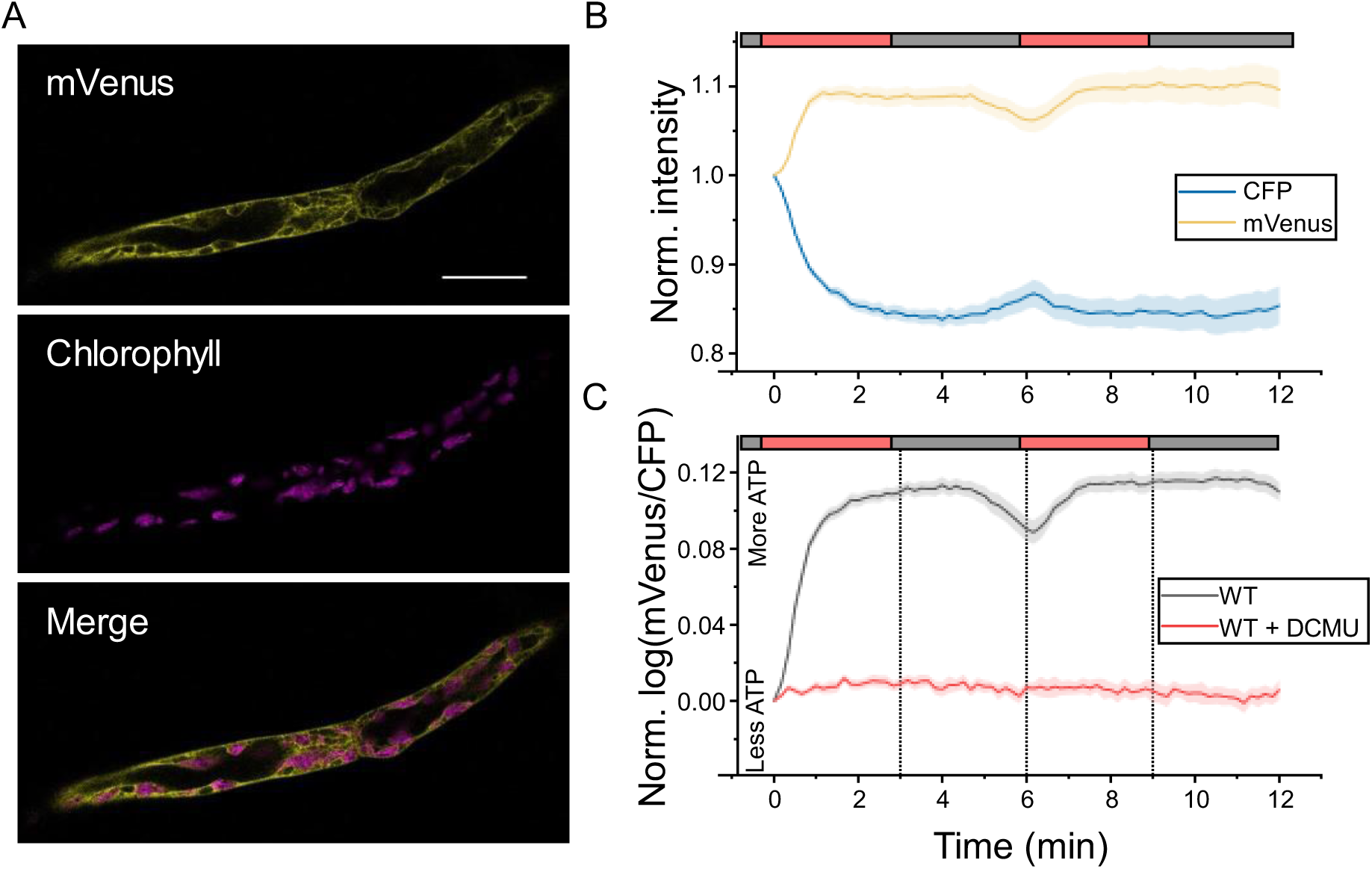
Light induced MgATP^2-^ accumulation in the cytosol of *Physcomitrium patens*. (A) Confocal images of two cells of protonema of one of the WT-ATeam lines showing mVenus channel (yellow), chlorophyll auto-fluorescence (magenta) and the merged image. Scale bar is 50 µm. (B) Normalized intensities of the single channels of WT-ATeam lines showing their changes upon dark-to-light transitions. Plants were dark adapted for 30 min before the measurements, and the first datapoint was measured before switching light on. Light periods are marked with a red bar and correspond to 50 μmol photons m^−2^ s^−1^ of red light. Error bands represent the SE of mean (n=40) (C) Normalized log-transformed FRET ratio during dark-to-light transitions of control and DCMU-treated WT-ATeam plants. Light periods are marked with a red bar and correspond to 50 μmol photons m^−2^ s^−1^ of red light. Error bands represent the SE of mean. Sample sizes are n=40 (WT) and n=13 (WT+DCMU). A live experiment is included as Supplemental Movie 1.

In the long term (> 1 year, corresponding to > 5 tissue regeneration) we observed a decrease in the fluorescence signal, most likely due to silencing effects, as similarly observed for different biosensor proteins in Arabidopsis (Schwarzländer et al., 2016; De Col et al., 2017) and for other overexpressed proteins in *P. patens* (Kubo et al., 2017). All imaging experiments were performed on protonema, a young tissue regenerated vegetatively, and we did not observe any mosaic silencing of the probe, meaning that the silencing, if present, was homogenous in all the cells. Because the probe is ratiometric, a moderate silencing would not cause any drift in FRET, as the value is self-normalized. In any case, if silencing effects appeared, new WT-ATeam plants were isolated to ensure maintenance of a good signal-to-noise ratio.

### Photosynthesis drives the increase of ATP concentration in the cytosol during dark-to-light transitions

In each round of transformation, three independent lines generated were used to observe the cytosolic MgATP^2-^ dynamics in the cytosol during dark-to-light transitions and assess changes in MgATP^2-^ levels associated with the activation of photosynthesis. To this end, photosynthesis activity was induced using 50 μmol photons m^−2^ s^−1^ of red (λ > 630 nm) actinic light. Red light is absorbed well by chlorophylls, but it does not interfere with the acquisition parameters of mVenus or CFP channels (465-561 nm). This choice of wavelengths thus enabled to keep the actinic light on during the acquisition of fluorescence without adding signal to the detector (Figure S1C, D). Taking advantage of the ratiometric nature of the probe and the natural distribution of protonema in single-cell layers, we quantified the CFP and mVenus signals and calculated the corresponding FRET ratio (mVenus/CFP) in whole focal planes that contained dozens of different cells (Figure 1A; Figure S1C).

To follow the dynamic response of ATP, plants were first incubated in the dark for 40 minutes to relax all photosynthesis-related processes and enable all samples to start from a homogeneous, dark-adapted state. When actinic illumination was switched on the mVenus signal increased immediately with a corresponding decrease in the CFP channel with the same kinetics (Figure 1B) a behaviour that indicates a *bona fide* increase in FRET efficiency between probes. mVenus/CFP ratio indeed increased within seconds of illumination, reaching a plateau after 1.5 minutes (Figure 1C). After light was turned off, FRET signal remained steady for approx. 1.5 minutes and then started to decrease, with a kinetic less steep than the light-driven increase (Figure 1C). When the light was switched on a second time, the signal increased again, confirming that MgATP^2-^ concentration responded to the light presence. In the second exposure to dark we observed a longer period of stability (about 2 minutes), before it started to decrease again.

The same measurements were repeated in the presence of the PSII inhibitor 3- (3,4-dichlorophenyl)-1,1-dimethylurea (DCMU) that blocks all photosynthetic electron transport, that completely abolished all the above-mentioned dynamics (Figure 1C; Figure S1E), demonstrating that light alone did not affect the FRET signal and that the observed dynamics were fully dependent on photosynthetic electron transport fuelled by the actinic light.

To assess the eventual effect of light quality on the cytosolic MgATP^2-^ dynamics, the same experiment was repeated using a different setup already used for similar experiments in Arabidopsis, where photosynthesis is induced with white light that is switched off during the confocal measurement, creating a pseudo-continuous light effect (Elsässer et al., 2020). These experiments showed very similar results, confirming that the dynamics observed were indeed the result of MgATP^2-^ accumulation in cytosol induced by light, independently from the setup and light wavelength (Figure S2A-D). The comparison further validates that both on-stage live illumination approaches as usable.

To test the effect of light intensity on the increase of MgATP^2-^ during dark-to-light transitions, we compared the kinetics of FRET upon illumination with different intensities, namely 5, 50 or 00 μmol photons m^−2^ s^−1^ of red light (Figure 2).

**Figure 2.**
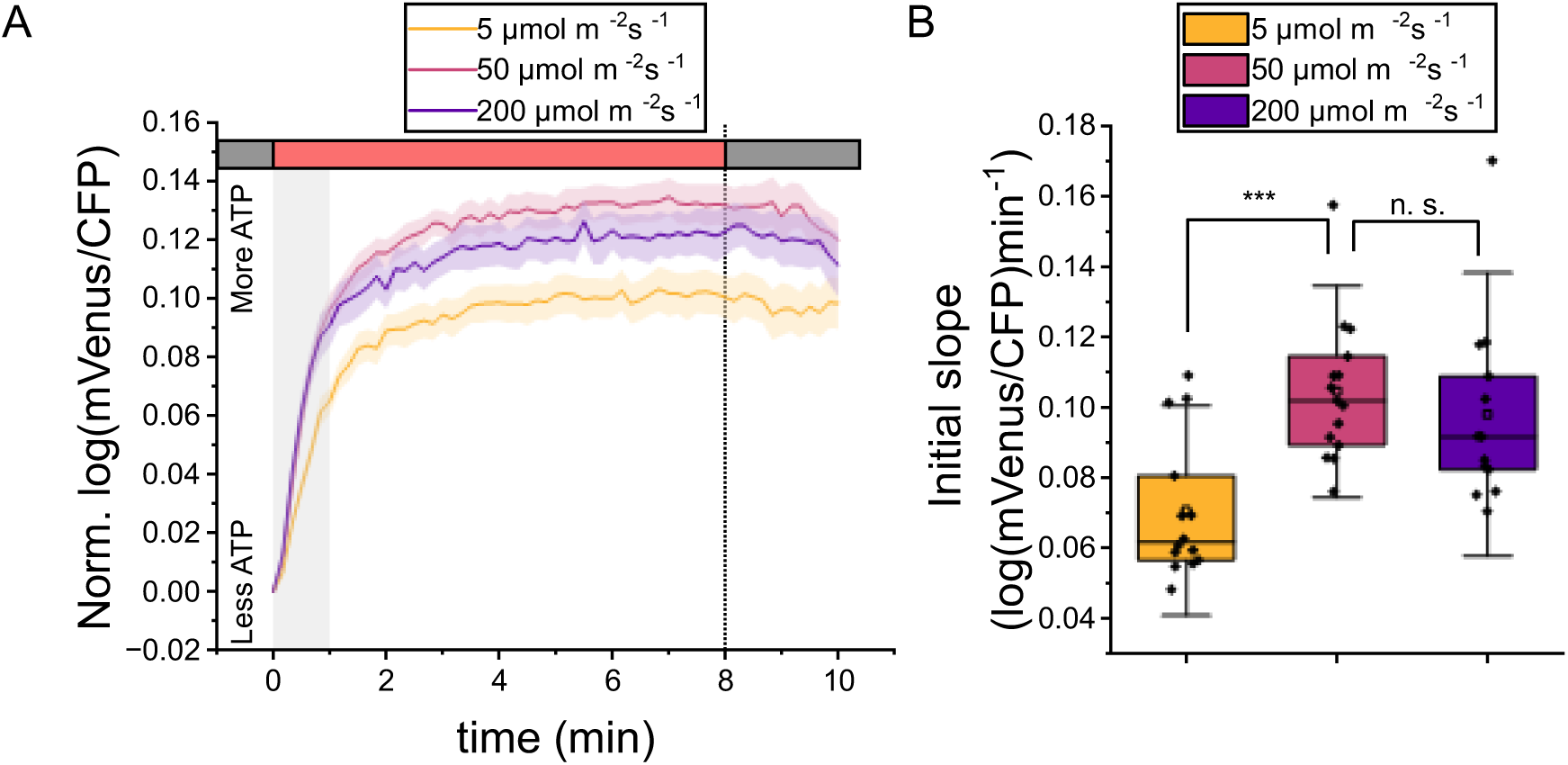
Effect of light intensity on the cytosolic increase of ATP during dark-to-light transitions. (A) Dark-adapted WT-ATeam plants were illuminated for 8 minutes with red light of either 5 (yellow , 50 (pink or 00 (purple μmol photons m^−2^ s^−1^, followed by 2 minutes of darkness. Dark and light periods are marked with grey or red bars, respectively. The shadowed region marks the timelapse used for the calculation of slopes shown in (B). The vertical dashed bar corresponds to the light-to-dark transition. Error bands represent the SE of mean. Sample sizes are n= (5 μmol photons m^−2^ s^−1^), n= 5 (50 μmol photons m^−2^ s^−1^ and n= 3 ( 00 μmol photons m^−2^ s^−1^). (B) Slope during the first minute of illumination. Statistics: two-sample t-test, (***) p<0.001; (n.s.) p>0.05. Error bars represent 1.5 times the SD.

An intensity of 5 μmol photons m^−2^ s^−1^ of red light, which is low and limiting for *P. patens* growth but high enough to initiate photosynthetic activity, was sufficient to trigger an increase in cytosolic MgATP^2-^ (Figure 2A), that was, however, slower than in 50 μmol photons m^−2^ s^−1^ (Figure 2B). This is in line with our finding that the increase in cytosolic ATP is fully dependent on the activation of electron transport in the thylakoids. On the other hand, using a higher light intensity ( 00 μmol photons m^−2^ s^−1^) did not alter the shape or magnitude of the increase of cytosolic ATP, even before reaching the plateau ( igure A, , meaning that 50 μmol photons m^−2^ s^−1^ of red light are sufficient to saturate ATP biosynthesis capacity in these conditions. This was also the case using the white light setup (Figure S2A).

### Mitochondrial respiration has seminal role in ATP supply in the cytosol

Mitochondrial activity has been suggested to contribute to cytosolic ATP biosynthesis in photosynthetic organisms and inhibitors of the mitochondrial respiratory chain have been shown to alter cytosolic ATP in Arabidopsis seedlings (De Col et al., 2017).

We monitored cytosolic ATP levels after blocking both the cyanide-sensitive and cyanide-insensitive electron transfer pathways, by treating plants with both KCN and salicylhydroxamic acid (SHAM), inhibitors of the Complex IV and the alternative oxidase (AOX), respectively. After this treatment, the basal ATP levels were drastically reduced (Figure S3C). Exposure to actinic light also triggered an increase of cytosolic ATP in KCN/SHAM-treated samples but with slower kinetics compared to control plants (Figure S2E), suggesting that mitochondrial respiration is a relevant contributor of cytosolic ATP biosynthesis. To assess impact of mitochondrial respiration on ATP biosynthesis using an orthogonal genetic approach, the ATeam1.03-nD/nA probe was introduced in *P. patens* lines lacking the Complex I structural subunit NDUFA5 (*ndufa5* KO) previously shown to completely lack mitochondrial NADH dehydrogenase activity (Mellon et al., 2021). Three independent lines stably expressing the cytosolic probe in *ndufa5* background, that we will refer to as *ndufa5*-ATeam, were isolated.

Complex I mutants showed impaired cell and tissue morphology: cells are smaller and protonema filaments more condensed than in WT (Mellon et al., 2021) (Figure S3A, B). The probe expression showed no additional impact on growth beyond the defects associated with *ndufa5* depletion (Figure S3A, B). The fluorescence intensity of the single channels was in all cases in a range close to the WT-ATeam lines and thus the same confocal settings were used for both lines to enable a reliable comparison of FRET signals between WT and *ndufa5* KO. There were no significant differences in the FRET ratio between WT-ATeam and *ndufa5*-ATeam dark-acclimated plants, meaning that the basal cytosolic ATP levels did not show significant differences between genotypes (Figure S3C).

*ndufa5*-ATeam plants were exposed to the same light treatment shown in Figure 1, observing an increase in ATP during dark-to-light transitions also in these mutants (Figure 3). However, the kinetics of FRET signal in *ndufa5*-ATeam plants was clearly affected and the initial slope 2.7 times slower than in WT-ATeam lines (Figure 3B). Differently to WT-ATeam, the signal also rapidly decreased when the light was turned off after the first 3 minutes of illumination. A second light treatment caused a further increase in cytosolic ATP, as in WT-ATeam, without reaching a plateau as observed in WT plants. These results suggest that mitochondrial respiration is an important contributor to the synthesis of cytosolic ATP and, remarkably, that Complex I is strictly required for efficient cytosolic ATP kinetics during dark-to-light transitions.

**Figure 3.**
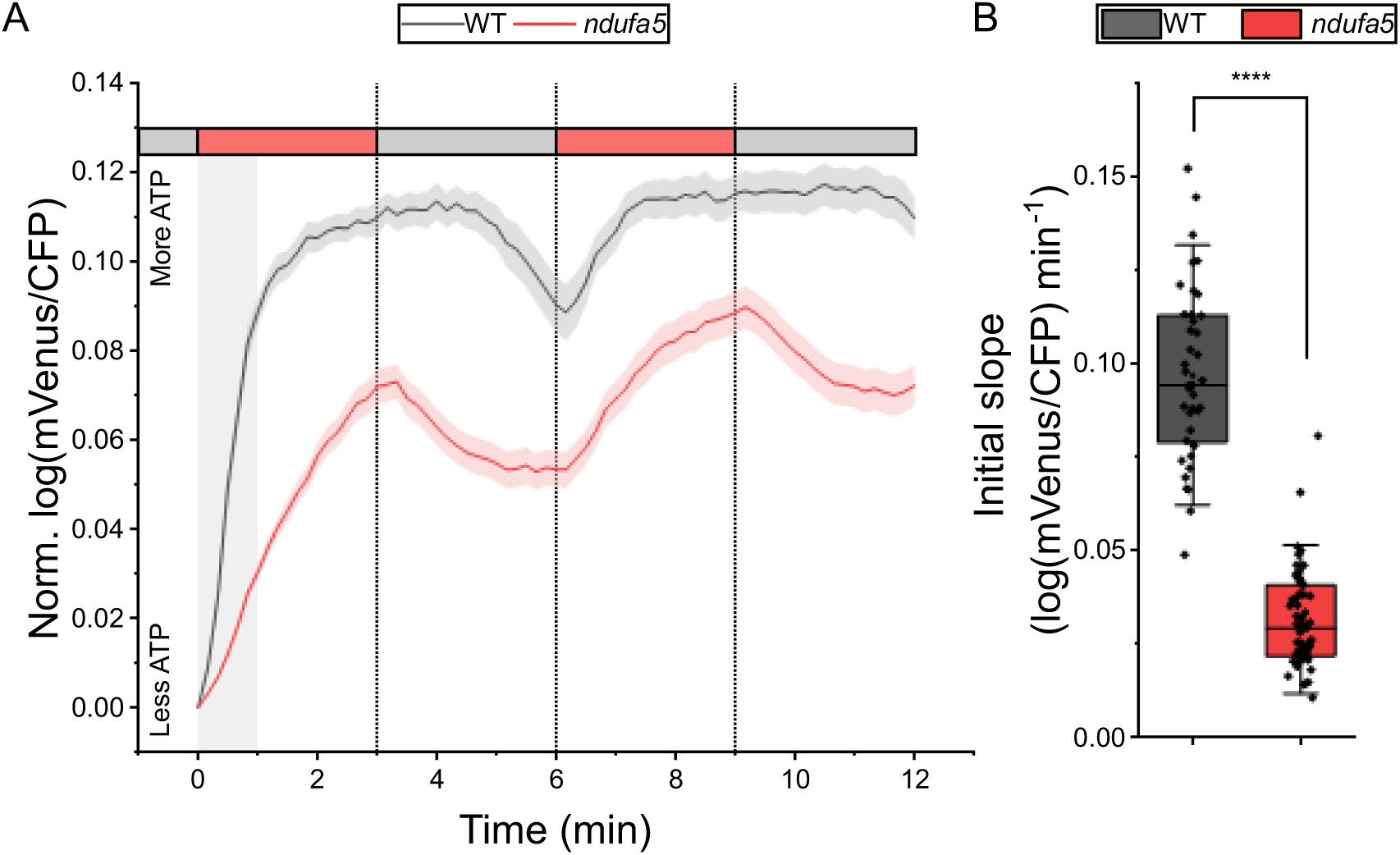
Cytosolic ATP dynamics in the Complex I deficient plant *ndufa5*-ATeam. (A) Normalized FRET ratio during dark-to-light transitions in WT and *ndufa5* KO. Dark and light periods are marked with grey or red bars, respectively. The shadowed region marks the timelapse used for the calculation of slopes shown in (B). The vertical dashed bars correspond to light-dark transitions. Error bands represent the SE of mean. Sample sizes are n=40 (WT) and n=55 (*ndufa5*). (B) Slope of the normalized FRET ratio during the first minute of illumination. Error bars represent 1.5 times the SD. Statistics: two-sample t-test, (****) p<0.0001.

To further investigate the ATP dynamics, WT-ATeam and *ndufa5*-ATeam plants were exposed to light treatments of different duration. The extension of the illumination phase up to 8 minutes (Figure 4A) showed that even though the rate of ATP biosynthesis was slower in the CI mutant, it almost reached the same normalized value of FRET as WT.

**Figure 4.**
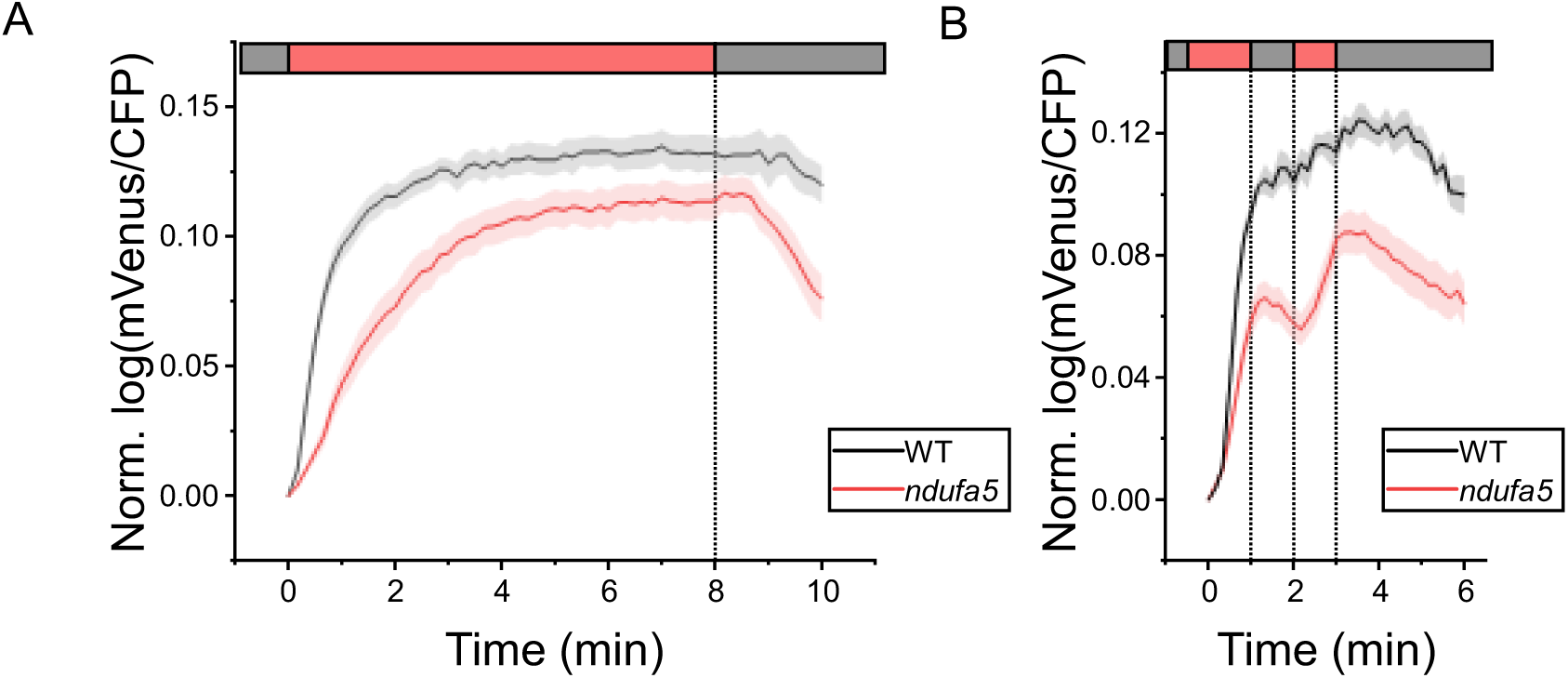
Cytosolic ATP dynamics in the Complex I deficient plant *ndufa5* under alternative light-dark cycles. Dark-adapted plants were exposed (A) to 8 min of light followed by 2 min of darkness or (B) to a light fluctuation that consisted in 1 min of light, 1 min of darkness, 1 min of light and 4 min of darkness. n both cases light used was 50 μmol photons m^−2^ s^−1^ of red light. Light and dark periods are marked with red and grey bars, respectively. Error bands represent the SE of mean. Sample sizes are n=15 (A, WT), n=15 (A, *ndufa5*), n=5 (B, WT) and n=25 (B, *ndufa5*).

On the other hand, when the illumination was reduced to 1 minute (Figure 4B), this time was sufficient for WT-ATeam to show a sustained increase in cytosolic ATP while one minute of darkness was instead not enough to observe a signal decrease. A following longer darkness period indeed showed that at least one and a half minutes of darkness were needed to observe the signal decrease. In *ndufa5*-ATeam, instead, FRET signal increase was slower during illumination, and it decreased immediately in the dark, confirming that rate of ATP biosynthesis was reduced in the mutant.

### Impairment of mitochondrial ATP biosynthesis drastically decrease growth without impacting photosynthesis

To further verify the biological relevance of mitochondrial ATP synthesis for plant metabolism we aimed to generate knock-out mutants of the mitochondrial F_1_F_O_-ATP synthase (complex V), the complex responsible of exploiting the electrochemical gradient generated by the mETC to synthesize ATP.

The F_1_F_O_-ATP synthase is largely conserved among eukaryotes with homologs of yeast and mammals subunits found conserved in green algae and plants and those could be identified in *P. patens* genome as well (Table S1). Proteomics approaches further identified two additional subunits in plants associated with the F_O_ domain, referred as F_A_d and 6 kDa, that present no counterparts in mammals or yeast (Senkler et al., 2017). Both subunits are also conserved in *P. patens* genome (Table S1). In particular, the subunit F_A_d has been linked with development and fertility in wheat where its repression leads to sterile plants(Li et al., 2010). In *Arabidopsis*, the gene is highly expressed in pollen during late developmental stages and the homozygous mutants are not viable. The hemizygous mutant shows altered mitochondrial morphology during the dehydration phase of pollen, causing their degeneration (Li et al., 2010).

Considering its functional impact and the fact that a single nuclear gene encodes for F_A_d, this subunit was chosen as target for the inactivation of complex V in *P. patens.* Two independent *f_A_d* lines, depleted of the gene Pp3c9_7910, were isolated and verified to have an insertion of the resistance cassette in the locus of interest (Figure S4A). This was possible because *P. patens* tissues are haploid in most developmental stages and the transformation procedure proceeds by vegetative propagation and it is thus possible to generate full knockout plants without passing through heterotrophic developmental stages, like spore formation and germination.

*f_A_d* plants showed a pronounced growth defect, much stronger than the one observed in the complex I deficient *ndufa5* ( igure 5A . mutants’ growth could not be rescued by exposure to continuous illumination or external feeding with glucose.

The pronounced nature of the growth defect limited the experimental analyses that were feasible for these plants. Yet, after approximately 4 months it was possible to obtain enough tissue to measure photosynthetic performance using chlorophyll fluorescence (Figure 5B). Upon exposition to mild illumination (50 µmol photons m^-2^ s^-1^), efficiency of photosystem (PS)I (Y_I_) was indistinguishable between WT and *f_A_d* plants. The efficiency of PSII (Y_II_) instead in the same conditions showed larger saturation and correspondingly the mutant showed a stronger reduction of plastoquinone, as estimated from 1-qL, suggesting that PSII electron transport capacity was reduced in the mutant with respect to the WT. The same measurements repeated with a stronger, sub-saturating light intensity (330 µmol photons m^-2^ s^-1^), however, showed instead no difference between WT and the Complex V mutant (Figure S4C), suggesting the photosynthetic light conversion is functional in the mutant. The mild alterations in photosynthetic activity observed could also be due to the different age and growth rate of plants but in any case, they cannot explain the major growth phenotype of the mutants.

**Figure 5.**
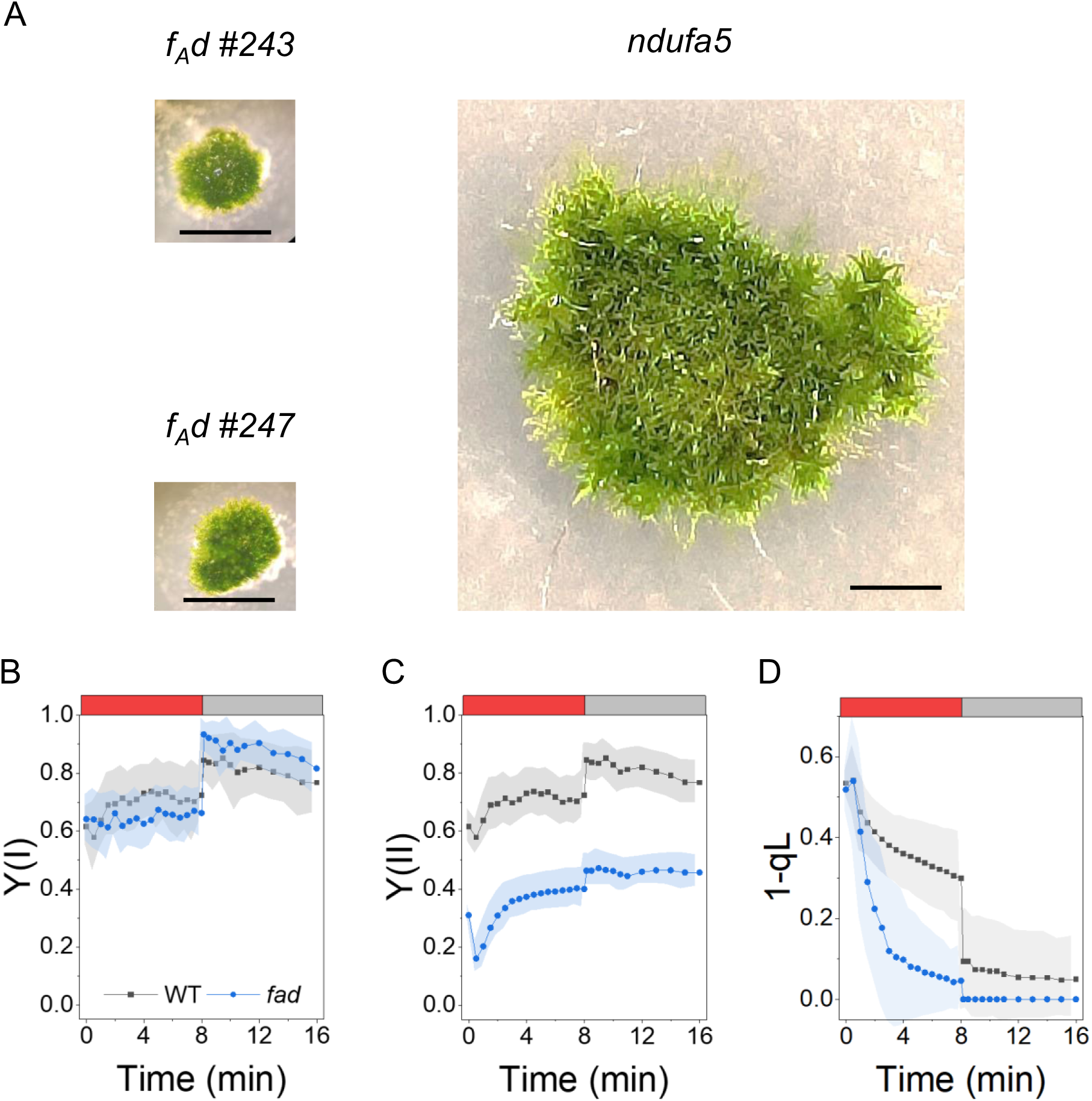
Growth phenotype and photosynthetic properties of *f_A_d* plants. (A) Comparison of 4 months-old *ndufa5* and *f_A_d* colonies. Scale bar is 2 mm. WT reaches the same size as *ndufa5* shown here in 2 weeks and cannot be cultivated for months because of depletion of nutrients in the medium (B, C) Quantum yield of Photosystem I (Y_I,_ B) and Photosystem II (Y_II_, C) monitored with PAM during exposition to 8 minutes of actinic light at 50 µmol photons m^-2^ s^-1^ followed by 8 minutes of dark. (D) Redox state of plastoquinone assessed by the fluorescence parameter 1-qL. All kinetics were measured after 40 minutes of dark adaptation. WT and *f_A_d* KO plants were grown photoautotrophically for respectively 10 days and 4 months. Data are expressed as the means ±SD, n = 4.

## Discussion

### Light driven ATP accumulation in the cytosol of P. patens

Dark-adapted cotyledons of Arabidopsis seedlings exhibited an illumination-driven increase in ATP levels (Voon et al., 2018). However, due to the high and saturating steady-state levels of cytosolic ATP in Arabidopsis cotyledons, light-induced differences were only observable when ATP levels were reduced before illumination through pre-treatment with the Complex I inhibitor rotenone, ensuring observations within the dynamic range of the probe (Voon et al., 2018). *Physcomitrium patens* lines expressing the FRET-based sensor ATeam1.03-nD/nA did not require pre-treatment with inhibitors. Their analysis revealed that plant illumination triggers a rise in cytosolic ATP in the absence of pharmacological inhibition of mitochondrial electron transport. This increase is entirely dependent on photosynthetic activity, as it is fully abolished by the PSII inhibitor DCMU. Using different light intensities, it was determined that 50 µmol photons m^-2^ s^-1^ of red light, a wavelength efficiently absorbed by chlorophylls, was already saturating for the experimental system employed here.

The MgATP^2-^, as the physiologically most relevant form of ATP, is stable in the slightly alkaline pH milieu of the cytosol leading the stable sensor readings between pH 7.5 and 8.5 (De Col et al., 2017). While illumination induces alkalinization in various compartments in Arabidopsis it was verified that it did not cause an increase in ATeam FRET (Elsässer et al., 2020). Hence, the most plausible explanation for the ATeam response in *P. patens* is a *bona fide* rise in ATP concentration. Under the assumption of no major changes in total adenylate pool size in the observed time frame, this indicates an increase in adenylate charge.

The differences in cytosolic ATP basal levels and dynamics between Arabidopsis and *P. patens* could be associated to the evolutionary distance between the species or to a different developmental stage, Arabidopsis seedlings vs. *P. patens* protonema cells in minimal medium and fully photosynthetically active. Remarkably, *P. patens* can have a life cycle completely photoautotroph, which is not the case for Arabidopsis.

In *P. patens* WT plants, the light-induced increase in FRET signal reaches a plateau approximately 1 minute after illumination. Conversely, it takes around 1.5 minutes of darkness for the FRET signal to decrease (Figure 1). This dynamic behaviour was highly reproducible. A potential technical explanation for the presence of a plateau may be probe saturation with MgATP^2-^, where the probe cannot resolve any further increases in MgATP^2-^, as previously observed in Arabidopsis(Voon et al., 2018). In that case MgATP^2-^ concentration surpasses the probe’s binding maximum of about 3 mM (at 25°C) (De Col et al., 2017) after 1 minute of illumination, and it takes 1.5 minutes in the dark to consume enough ATP to re-enter the probe’s dynamic range. If this was the case, however, in experiments with shorter illumination (1 minute) FRET signal would be expected to decrease immediately after the light is switched off. The ATeam signal remains stable after the end of the illumination phase for approximately 1.5 minutes, independent of the duration of the light treatment (1, 3, or 8 minutes). An alternative biological explanation consistent with all observations is instead that MgATP^2-^ concentration reaches a steady state as a result of ATP buffering by a characteristic feature of the metabolic system and/or its regulation.

This ATP buffering activity can be attributed to mitochondrial respiration that utilizes reducing equivalents produced upstream by photosynthesis and transported to the mitochondria through redox shuttles (Igamberdiev and Bykova, 2022), thus maintaining a supply of ATP in the cytosol. Following this hypothesis the decrease of reducing equivalent export from the chloroplast would thus only start impacting cytosolic ATP levels if the dark period exceeded 1.5 minutes. The concept of ‘metabolic lag’ between chloroplasts and mitochondria is not new, as it is the most accepted hypothesis to explain the phenomenon referred to as light-enhanced dark respiration (LEDR) (Lehmann et al., 2016). This phenomenon describes the large value of respiration that can be measured, either as O_2_ consumption or CO_2_ evolution, in plants immediately after exposure to dark. Values of respiration will then approach a lower steady value in circa 30 minutes (Azcón-Bieto and Osmond, 1983). Behind LEDR it is believed to be the accumulation of reduced substrates, exported by chloroplasts, that are still available for mitochondria for some minutes even though they are no longer produced after a recent light exposure (Gessler et al., 2017). These can drive respiration, which could as well be responsible for the ‘residual’ ATP production that we describe here as ATP buffering.

Another important conclusion that can be drawn from the observation of rapid cytosolic ATP accumulation after onset of illumination and the delayed decrease after the onset of darkness is an ability of the plant cells to mitigate the metabolic impact of changes in illumination intensity, maintaining relatively steady ATP levels even in the presence of abrupt light fluctuations. In a natural and dynamic environment, light changes are common and significantly impact photosynthetic productivity and yield in crops (Wang et al., 2020; De Souza et al., 2022). The observed kinetics imply that while photosynthetic activity responds rapidly to changes in light availability, plant cells can stabilize their cytosolic ATP levels to uncouple cytosolic energy physiology from abrupt light fluctuations, while maintaining a link to long-lived changes in illumination. Using mitochondrial respiration as an integrated strategy to supply the cytosol with photosynthesis-derived ATP, rather than taking a more direct strategy such as the triose phosphate shuttle, may provide a decisive advantage under natural highly dynamic circumstances. As such the ability to maintain photosynthesis-derived ATP supply of the cytosol stable by taking the ‘detour’ via mitochondrial respiration may represent a central element of the strategy of plants to cope with dynamic light conditions and to thrive under natural growth conditions (Long et al., 2022).

### Mitochondrial respiration is essential for cytosolic ATP accumulation

About one minute of illumination is sufficient to elevate cytosolic MgATP^2-^ levels to a steady state in wild type. That steady state is maintained for 1.5 minutes after onset of darkness (Figure 4B). Contrarily, in *ndufa5* plants lacking functional mitochondrial complex I, MgATP^2-^ accumulation in the cytosol occurs at an approximately three times slower rate. As a result of slower accumulation, a steady-state in ATP levels is reached only after prolonged light exposure of up to 8 minutes. Then similar MgATP^2-^ levels are reached as in wild type.

The impact of respiration in cytosolic ATP supply is confirmed by the dynamics immediately after light cessation. In the *ndufa5* background, the sensor FRET signal decreases promptly in the dark, whereas in WT background, it remains stable for approximately 1.5 minutes. This response behaviour supports the hypothesis that mitochondrial respiration contributes to ATP supply to the cytosol not only during illumination but also during dark-to-light transitions. *ndufa5* plants, deficient in Complex I, which contributes to 40 % of proton translocation for mitochondrial ATP biosynthesis (Braun et al., 2014), exhibit a significant reduction in ATP accumulation in the cytosol. The degree by which *ndufa5* plants are affected, even though the lines retain proton translocation capacity from respiratory complexes III and IV, emphasizes the importance of functional mitochondrial respiration to act as major contributor to cytosolic ATP supply in plants.

The essential role of mitochondrial ATP biosynthesis is further underscored by the phenotype of mutants (*F_A_d*) with impaired mitochondrial ATP biosynthesis, showing massive growth reduction even under continuous illumination. The observation that those mutants exhibit only mild alterations in photosynthetic activity, further emphasize that it is mitochondrial ATP biosynthesis and ATP supply to the cytosol that make respiration indispensable in photosynthetically active cells.

While the mutant data support that the mitochondrial contribution is crucial, all observed dynamics strictly depend on photosynthetic activity, as evidenced by light dependence and the abolition of ATP accumulation by the PSII inhibitor DCMU. This aligns with the hypothesis that chloroplasts predominantly export reducing equivalents, readily imported and utilized by mitochondria for ATP synthesis, which is promptly exported to the cytosol.

These findings provide critical and hitherto lacking *in vivo* validation of biochemical studies on plant cell ATP compartmentation, highlighting limited capacity of chloroplasts to export ATP in mature leaves. Our findings are consistent with the idea that mitochondria play a quantitatively major role in ATP supply to the cytosol, as proposed by previous work based on biochemical considerations (Millar et al., 2011; Gardeström and Igamberdiev, 2016). While alternative pathways for ATP supply may exist and are not ruled out, their inability to quantitatively compensate for mitochondrial inactivation, as observed in *f_A_d* mutants, emphasizes the pivotal and dominant role of mitochondria in cytosol ATP supply.

The use of *P. patens*, which is a bryophyte, allows us to elaborate further conclusions. Opposite to Arabidopsis, this model is photoautotrophic throughout its development. Therefore, the impact of depleting respiration is not due to presence of heterotrophic tissues that rely exclusively on respiration for obtaining energy and thus enable to conclude instead that the respiration is essential in plants for the supply of ATP in the cytosol in active photosynthetic tissues.

### Evolutionary perspectives on photosynthesis-dependent ATP biosynthesis in the mitochondria

Recent studies have shed light on the role of respiration in various photosynthetic organisms, including diatoms, green algae, and plants. While these investigations share a common recognition of the biological relevance of respiration, mechanistic differences have been observed. In the green alga *Chlamydomonas reinhardtii* photosynthesis and growth can proceed in photoautotrophic conditions even if respiratory Complexes III and IV are inactive(Salinas et al., 2014). This contrasts with plants, where an impairment of Complexes III and IV has a major impact on growth and their complete inactivation is lethal. This divergence can be attributed to the fact that *Chlamydomonas* can grow both autotrophically and heterotrophically. In its natural environment, it may encounter anoxic conditions, resulting in respiratory inhibition. In contrast, plants are mostly obligatory autotrophs and have adapted to the constant presence of oxygen, rendering them hypoxia sensitive(Loreti and Perata, 2020). Since plants are normally exposed to oxygen, respiration can be continually active in plant cells without running any major risk of dysfunction. That may be sufficient to account for the evolutionary adaptation of their photosynthetic energy metabolism to strictly work in concert with respiration.

Delegating cytosolic ATP supply to mitochondrial respiration in plant cells generates a futile cycle under illumination where O_2_ and reducing power generated by photosynthesis are consumed by mitochondrial respiration. This allocation of metabolic roles between mitochondria and chloroplasts must thus generate another evolutionary advantage where the benefits of relying on mitochondrial respiration for cytosolic ATP supply outweigh the efficiency cost.

One possible advantage that is suggested by our observations is the ability to maintain a steady ATP supply to the cytosol. Buffering photosynthesis-derived ATP supply to the cytosol under changeable light conditions may help mitigating changes in photosynthetic activity induced by rapid light fluctuations while maintaining a steady supply of ATP to maintain critical housekeeping functions. In contrast, if ATP supply were more directly linked to photosynthetic activity, cytosolic ATP availability would closely follow illumination dynamics, resulting in rapid dynamics in concentration (and possibly energy charge) following any alteration in light availability.

Another potential advantage may lie in the regulation of photosynthesis. The proton gradient (ΔpH across the thylakoid membrane serves as the energy source for ATP biosynthesis in the chloroplast but also is a key signal for the regulation of photosynthesis, controlling the modulation of multiple mechanisms such as nonphotochemical quenching (NPQ), xanthophyll cycle and photosynthetic control(Eberhard et al., 2008). If mitochondria are the primary contributors to ATP supply for the cytosol, this will allow the metabolic demand for ATP supply to be decoupled from photosynthetic regulation. In an alternative scenario where ATP synthesis is localized in the chloroplast, limiting ATP synthase activity to increase ΔpH and induce photosynthesis regulation would also restrict ATP supply to the cell. Under stress conditions, active regulation of photosynthesis via regulating the activity state of ATP-synthase may thus impede ATP supply to other parts of the cell, potentially impairing essential functions of housekeeping and the ability to mount efficient stress responses, such as membrane energization, active transport, cytoskeletal remodelling and gene expression.

In both scenarios, photosynthesis-derived ATP supply via mitochondrial respiration enables plants to respond effectively to variable environmental conditions. This adaptability is a decisive evolutionary driving force for plants and may even outweigh potential disadvantages in simple energy efficiency. Hence, the data we present here offer in vivo evidence that mitochondrial respiration is strictly required and maintained in plants to ensure cellular supply with photosynthesis-derived ATP.

## Methods

### Plant materials and growth conditions

The wild-type strain used in this study was the Gransden wild-type (WT) strain of *Physcomitrium patens* subsp. *patens*. Two different media were used for growing protonemal tissue, the minimal PpNO_3_ medium and the enriched version PpNH_4_(Ashton et al., 1979). PpNO_3_ medium contained, per litre, 250 mg MgSO_4_ · 7 H_2_O, 800 mg Ca(NO_3_)_2_ · 4 H_2_O, 12.5 mg FeSO_4_ · 7 H_2_O, 1 mL KH_2_PO_4_-KOH buffer and 1 mL trace element solution. KH_2_PO_4_-KOH buffer contained 25 g KH_2_PO_4_ per 100 mL and pH 7 was obtained by titration with 4 M KOH. Trace element solution contained 55 mg CuSO_4_ · 5 H_2_O, 55 mg ZnSO_4_ · 7 H_2_O, 614 mg H_3_BO_3_, 389 mg MnCl_2_ · 4 H_2_O, 55 mg CoCl_2_ · 6 H_2_O, 28 mg KI, 25 mg Na_2_MoO_4_ · 4 H_2_O per litre. Medium was solidified with 8 g/L agar and sterilized by autoclaving. PpNH_4_ medium resulted from supplementing PpNO_3_ medium with 500 mg ammonium tartrate and 5 g glucose per litre. Solidification of PpNH_4_ medium was achieved with 7.2 g/L agar. The final pH of sterilized media was 5.5 to 6.0.

For propagation, protonemal tissue was grown on cellophane-layered PpNH_4_ medium at 24°C under moderate light intensity (50– 0 μmol photons m^-2^ s^-1^) and long day conditions (16 h light, 8 h darkness) in growth chambers. WT and WT-ATeam plants were harvested after 5 or 6 days of growth, and *ndufa5* and *ndufa5*-ATeam plants were harvested after 7 to 10 days. *f_A_d* plants were grown for months. Growth of *f_A_d* KO lines was also tested under continuous illumination. In this case, temperature and light conditions were unchanged, with removal of photoperiod. For confocal imaging experiments, protonemal tissue was grown on cellophane-layered minimal PpNO_3_ medium under the same conditions and used after 10 days of growth.

### Moss transformation and transgenic plant selection

For generation of ATP reporter lines, the *P. patens* cytATeam_pT1OG expression vector was generated as follows. The coding sequence of ATeam1.03-nD/nA flanked by the Gateway *ntt* regions was isolated from the Arabidopsis Gateway expression vector pH2GW7_cytATeam1.03.nDnA(De Col et al., 2017) using the enzyme SbfI. The Gateway BP reaction was done using the previous linearized fragment and the donor vector pDONR221. After BP reaction, the entry clone pDONR221-ATeam1.03.nD.nA was used to transform DH5α *E. coli* thermocompetent cells. The kanamycin resistant colonies were screened by PCR using M13for and M13rev primers, and positive colonies used for minipreps. Identity of pDONR221-ATeam1.03.nD.nA minipreps was verified by restriction with BsrGI or EcoRV+ApaI. The Gateway LR reaction was done using the entry clone and the destination vector PT1OG (Aoyama et al., 2012, GenBank ID LC126301). After LR reaction, the expression vector cytATeam_pT1OG was used to transform DH5α *E. coli* thermocompetent cells. Some of the kanamycin resistant colonies were used for miniprep. The identity of the expression vector cytATeam_pT1OG was verified in the minipreps by restriction with BsrGI or PmeI.

Transformation of *P. patens* was carried out by the polyethylene glycol (PEG)-mediated method (Nishiyama et al., 2000), using either protoplasts of WT or *ndufa5* lines (the latter published previously(Mellon et al., 2021)) to generate reporter lines in both WT and *ndufa5* backgrounds. Protoplasts prepared from one-week-old *P. patens* protonemata were incubated with 2 % (w/v) Driselase in 8.5 % (w/v) mannitol for 30 minutes. Protoplasts were washed three times with 8.5 % (w/v) mannitol and suspended at ∼1.6 × 10^6^ cell/mL in a solution containing 8.5 % (w/v) mannitol, 15 mM MgCl_2_ and 0.1 % (v/v) MES (pH 5.6). Linearized DNA of cytATeam_pT1OG (20 µg) was mixed with the protoplast suspension (300 μL and the PEG solution (300 μL containing 7 % (w/v) mannitol, 40 % (w/v) PEG4000, 100 mM Ca(NO_3_)_2_ and 10 mM Tris (pH 7.2), and incubated at 45°C for 5 minutes in a water-bath. After 10 minutes of incubation at room temperature, protoplasts were suspended in liquid protoplast regeneration medium (6.5 mL) obtained by supplementation of liquid PpNH_4_ with 6.6 % (w/v) mannitol and incubated overnight at 24°C in the dark condition. 7 mL of melt top layer solution, containing 8.5 % (w/v) mannitol and 14 g/L agar, were poured onto the protoplast-containing mixture at approximately 50 °C and mixed gently. The resulting mixture was spread over four cellophane-overlayed Petri dishes (3.5 mL each) containing solid protoplast regeneration medium. After 6 days of incubation at 24 °C under light condition, the regenerated cells were transferred onto a PpNH_4_ agar medium containing the antibiotic zeocin (50 ng μL . Antibiotic resistant colonies were transferred and grown for 7 to 14 days on antibiotic free PpNH_4_ medium, and then transferred again to antibiotic containing PpNH_4_. Screening of stably resistant lines was done through quantification of mVenus fluorescence, as described in the main results section.

For generation of plants missing a functional mitochondrial ATP synthase, we generated the Fad_KO_BHRf construct by inserting the PCR-amplified flanking regions of the locus encoding F_A_d (Pp3c9_7910) (Figure S4A) in the BHRf vector (kindly provided by F. Nogue, INRA Versailles, France). Transformation of *P. patens* was carried out as described for ATP reporter lines. In this case the antibiotic used for screening was hygromycin (5 ng μL and screening of stable resistant lines was done through PCR verification of disruption of the genomic F_A_d locus (Figure S4B). Genomic DNA extraction protocol was adapted from a previously published protocol (Edwards et al., 1991) and was performed as follows. A piece of stable resistant colony was collected in a 2 mL screw-cap tube containing three 3-mm zirconium glass beads. After addition of 500 μL of cold TE buffer ( 00 mM Tris-HCl, pH 8.0; 50 mM EDTA; 500 mM NaCl), tissue was homogenized through two consecutive cycles of bead beater at maximum speed for 30 seconds, with 1 minute of cold incubation between cycles. After the addition of 35 μL of 0 % (w v S S, samples were incubated at 65 ° for 5 minutes. Then, 30 μL of potassium acetate 5 M were added, samples were kept on ice for 5 minutes and centrifuged at 4 °C for 10 minutes at 13,000 g. Supernatant was transferred to a clean 1.5 mL tube containing 500 μL of isopropanol at -20 °C, mixed by inversion and incubated at -20 °C for 10 minutes. Then a series of centrifuges at 4 °C and 13,000 g were performed. After a first centrifuge of 10 minutes, the supernatant was discarded, and the pellet washed with 500 μL of 70 % ethanol. Then, the resultant pellet of a second centrifuge of 5 minutes was washed with 50 μL of 70 % ethanol. A last centrifuge of minutes was performed, and the resultant pellet was dried off under a fume hood. The pellet was then resuspended in 50 μL of purified water and this A solution was kept at -20 °C until used. PCR amplifications of the recombination cassette were performed on extracted gDNA. Primers used for construct design and line validation are the following (5’ to 3’ sequence . Fad-p1: GTGGAATGGCATGATTTTAT; Fad-p2: ACTAAACAACCAGACCAGGA; Fad-p3: CTTGGGATTGATGATGCTAT; Fad-p4: TGGGTATTATCGGAGTCAAC; Fad-p5: CATCTACCATTTTGGGTTTC; Fad-p6: ACTTCGAACCAGTTCCAGTA; HNZ Up Rev: TGCGCAACTGTTGGGAAG; 35S terminator: CGCTGAAATCACCAGTCTCTCT.

### Confocal imaging

Plants were grown for 10 days on solid PpNO_3_ medium and dark adapted for 40 min. A piece of protonema was extended on a slide with a drop of liquid PpNO_3_ medium (without agar) and covered with a coverslip. The coverslip was then fixed with duct tape to avoid desiccation of the sample during the measurement. This process was done in a dark room to keep the samples dark-adapted.

Samples were illuminated with 50 μmol photons m^−2^ s^−1^ using a red-filtered (580-630 nm) light (halogen lamp connected to an optic fibre). Light intensity was set to 50 μmol photons m^−2^ s^−1^ using a light meter (LI-COR LI-250A). The spectrum of the filtered light is included in the Figure S1D and was measured using a LI-COR LI-180 Spectrometer.

Imaging was performed using a 40× oil immersion lens using a Leica SP5 confocal microscope (Leica Microsystems). An Argon laser (488 nm) was used to excite both CFP and chlorophylls, with laser power set at 12.5 %. A scan was performed every 10 seconds. The first scan was done at darkness to quantify the basal ATP levels. Light was then turned on and off according to the described protocols. Three emission channels were set up: CFP (465-500 nm), mVenus (525-561 nm) and auto-fluorescence of chlorophylls (670-690 nm) (see Figure S1).

The acquired fluorescence datasets were analysed using the MatLab-based Redox Ratio Analysis software (RRA) (Fricker, 2016). For each dataset, five square regions of interest (ROIs) were analysed. Each ROI was analysed as a single replicate. First, background signal was removed using the integrated functionality of the RRA software. The intensities of the single mVenus and CFP channels were then exported and used for further analysis and plotting with OriginPro. We calculated the mVenus/CFP intensity ratio, which was then log-transformed to approach the normal distribution. The log-normalized values were then normalized to the value at time 0, i.e., to the steady state in the dark.

The slope of the increase in the ATeam signal during dark-to-light transitions was calculated by fitting the datapoints (norm. log(mVenus/CFP)) to the formula y=a+bx using the Linear Fitting tool of OriginPro, where b is the reported slope. Statistical significance between two samples was analysed by two-tailed Student’s t test (alpha = 0.05) using OriginPro. The number of replicates and meaning of asterisks are indicated in the figure legends.

### Chlorophyll fluorescence experiments

Photosynthetic parameters were retrieved from chlorophyll fluorescence to analyse photosystem II (Y_II_, qL) and a dual wavelength absorbance spectrometry to analyse photosystem I (Y_I_), as described previously (Gerotto et al., 2016). Chlorophyll fluorescence and near-infrared (NIR) absorption analyses were performed at room temperature using a Dual-PAM 100 system (Walz) on protonema grown for 10 days in PpNO_3_ in WT and for approximately 4 months for the fad mutant. Before the analysis, plants were adapted in the dark for 40 minutes. Induction curves were obtained by setting actinic red light at (approx.) 50 or 330 µmol photons m^-2^ s^-1^, and photosynthetic parameters were recorded every 30 seconds. At each step, the photosynthetic parameters were calculated as follows: Y_II_ as (F_m_’-F_o_)/F_m_’; qL as ( _m_’-F)/(F_m_’-F_o_’ x F_o_’ ; Y_I_ as 1-Y_ND_-Y_NA_, with Y_NA_ as (P_m_-P_m_’)/P_m_ and Y_ND_ as (P - P_o_/P_m_) (Klughammer and Schreiber, 1994).

## Author contributions

Conceptualization, A.A. and T.M.; Investigation, A.M.V.V., P.N., K.Z. and S.L.T.; Resources, M.S.; Supervision, M.S., A.A., T.M.; Writing – original draft, A.M.V.V and T.M.; Writing – review & editing, A.A, K.Z. and M.S.

## Acknowledgments

A.M.V.V. acknowledges the support from University of Padova and the Italian Society of Plant Biology. K.Z. and S.L.T. are grateful for support by the Chinese Scholarship Council. M.S. thanks the Deutsche Forschungsgemeinschaft (DFG) for funding though the infrastructure grant INST211/903-1 FUGG, and the project grants SCHW1719/5-3, SCHW1719/10-1 and SCHW1719/11-1. Authors thank Francesca Papini (University of Padova) for preliminary experiments.

