## Supplemental Material for "Mitochondrial respiration is essential for photosynthesis-dependent ATP supply of the plant cytosol"

### Supplemental information

**Supplemental Movie 1 |** Increase in FRET signal on a WT-ATeam plant during dark-to-light transitions. Left, live merge image of CFP emission, mVenus emission and brightfield channels. Right, plotted data taken from experiment shown at right. Related to Figure 1.

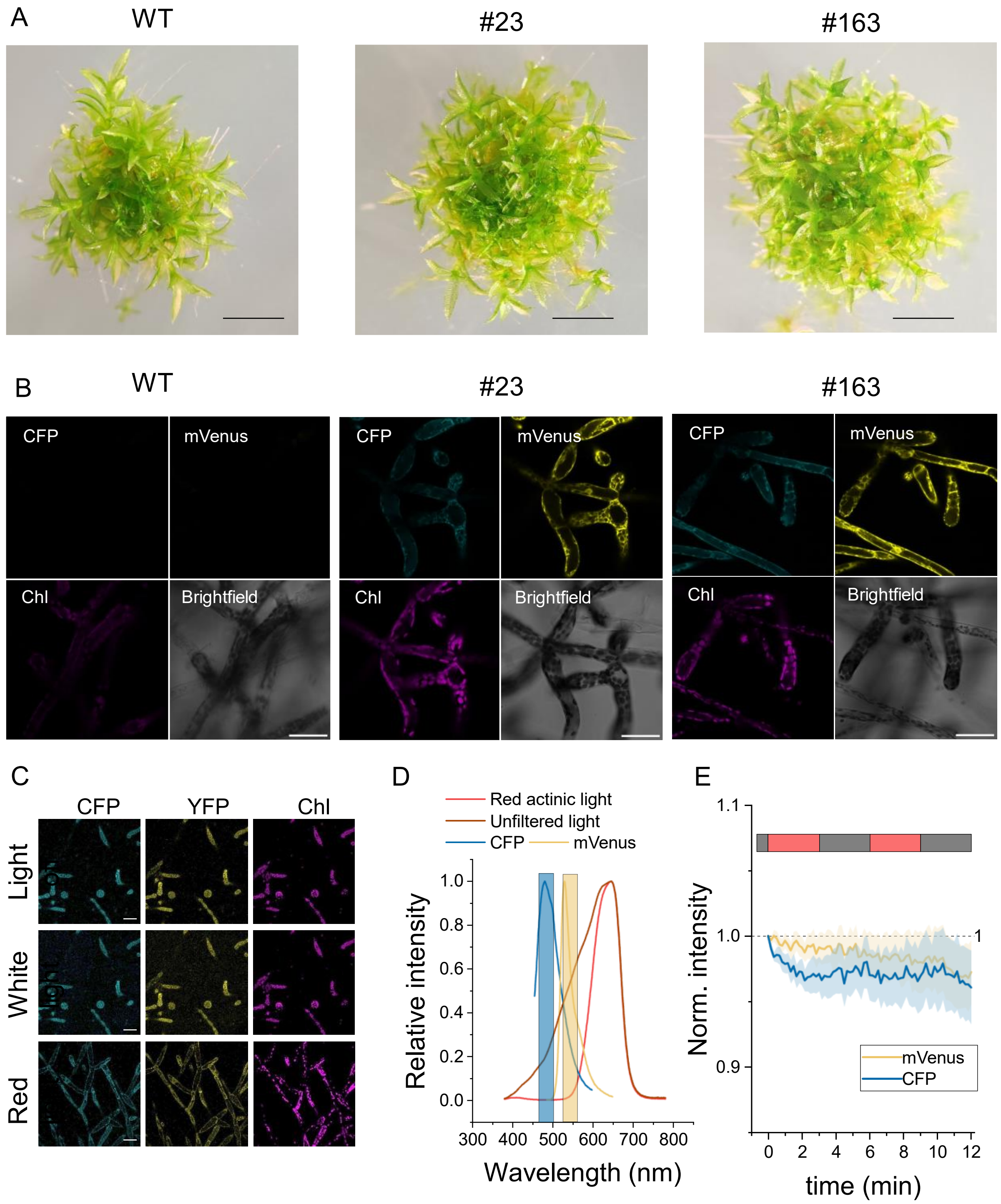


Figure S1 | **Isolation of WT-ATeam *P. patens* plants and verification of impact of actinic light on probes fluorescence determination.** As examples, two representative WT-ATeam lines, #23 and #163, are shown. (A) Plants after 28 days of growth in solid $PpNO_3$ minimal medium. Scale bars are 2 mm. (B) Fluorescence signal and brightfield images of 10 days old protonema. Scale bars are 50 µm. (C) White light does interfere with CFP and mVenus channels, as shown by the noisy background under white illumination, that is not present in the dark or when light is red. Scale bar is 50 µm. (D) Emission spectra of CFP (blue) and mVenus (yellow) and the corresponding acquisition ranges of the microscope (rectangles). Spectra of the actinic light used to induce photosynthesis before and after filtering are shown. (E) Signals of the single CFP and mVenus channels during dark-to-light transitions in DMCU treated plants. Light periods are marked with a red bar and correspond to 50 µmol photons $m^{-2}$ $s^{-1}$ of red light. Error bands represent the SE of mean (n=12).

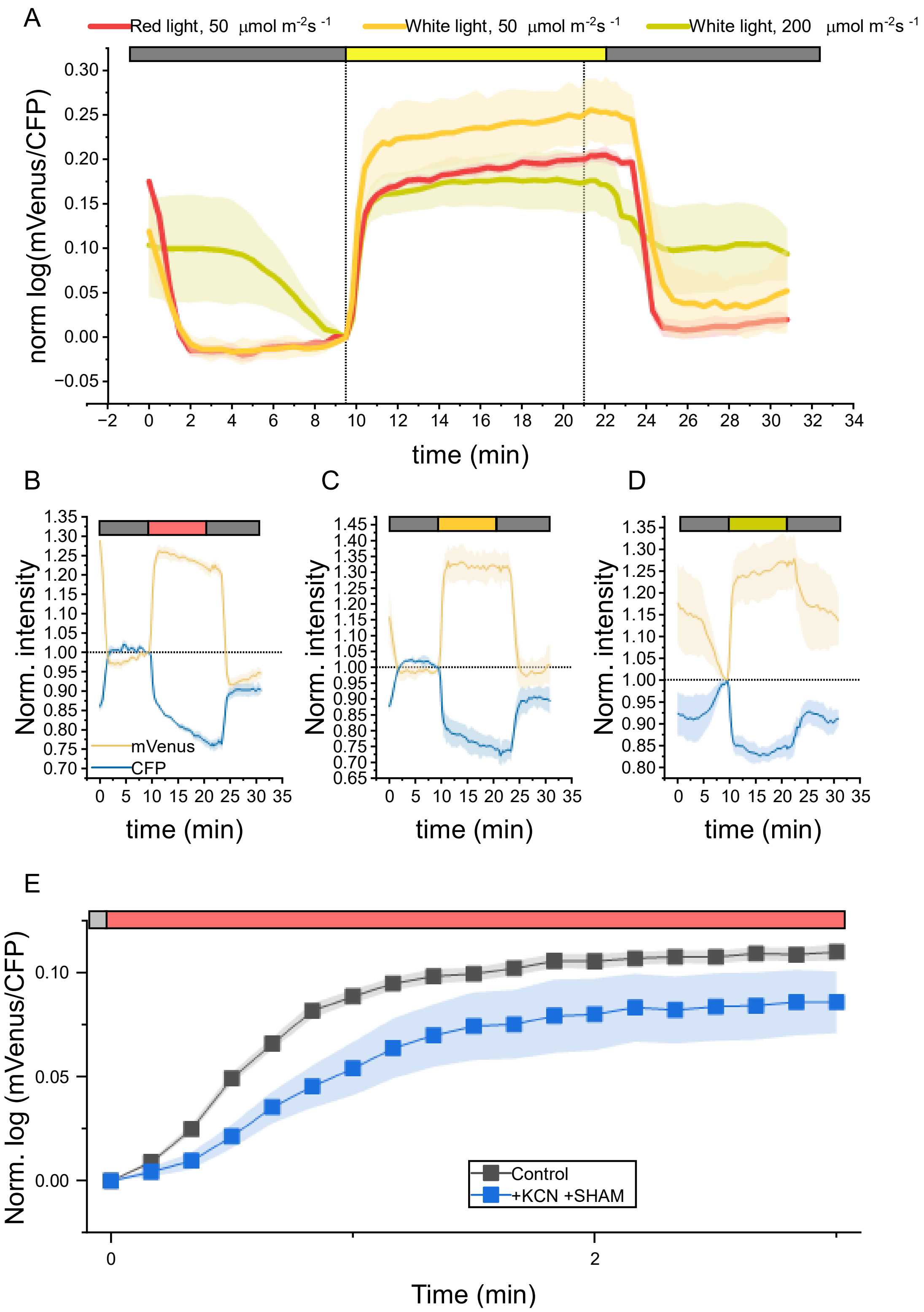


Figure S2 **| Cytosolic ATP dynamics during dark-to-light transitions using a pseudo-continuous light system as described in Elsässer et al 2020 [1]. or after blockade of respiration.** The mVenus/CFP ratio (A) was calculated for 10 minutes at darkness, followed by 12 minutes of illumination and further 10 minutes of darkness. We compared the effect of different light qualities (red or white light) and intensities (50 or 200 µmol photons $m^{-2}$ $s^{-1}$). The intensities of single CFP and mVenus channels are show in B-D. (E) Increase in cytosolic ATP during dark-to-light transitions of plants treated simultaneously with KCN and SHAM.

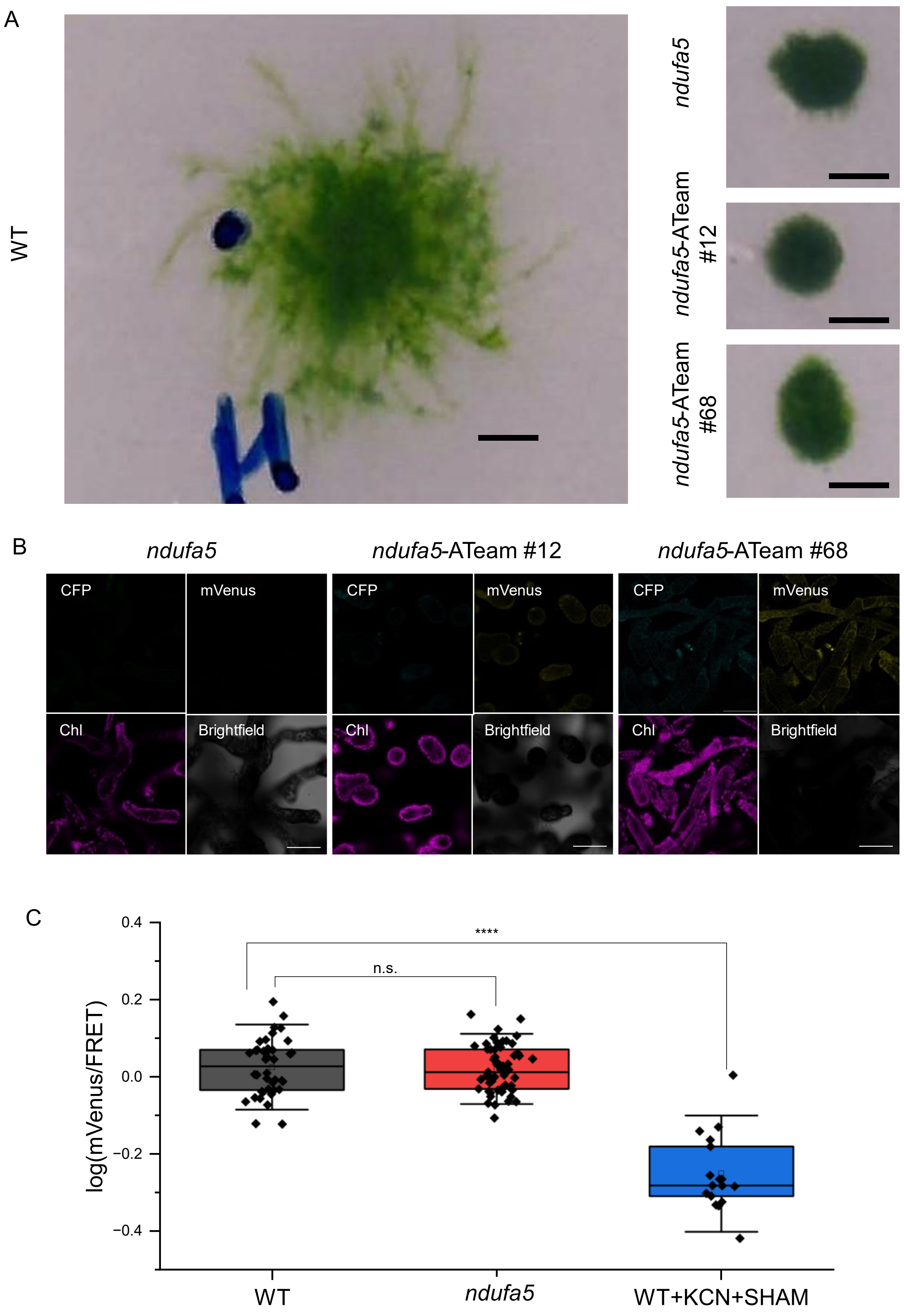


Figure S3 | **Isolation of *ndufa5*-Ateam plants and basal levels of cytosolic ATP in dark-adapted plants.** (A) Picture of colonies after 21 days of growth, showing parental *ndufa5* KO plant, two *ndufa5*-ATeam lines and a WT for comparison. All pictures are shown with the same scale, scale bar is 2 mm. (B) Fluorescence signal and brightfield images of 10 days old protonema. Scale bars are 50 µm. (C) Log-transformed mVenus/CFP ratio values of dark-adapted plants not exposed to actinic light. Inhibition of the respiratory chain by treatment with KCN and SHAM caused a strong decrease of basal cytosolic ATP levels. Statistics: two-sample t-test, (****) $p<0.0001$, (n. s.) non-significant. Error bars represent 1.5 times the SD.

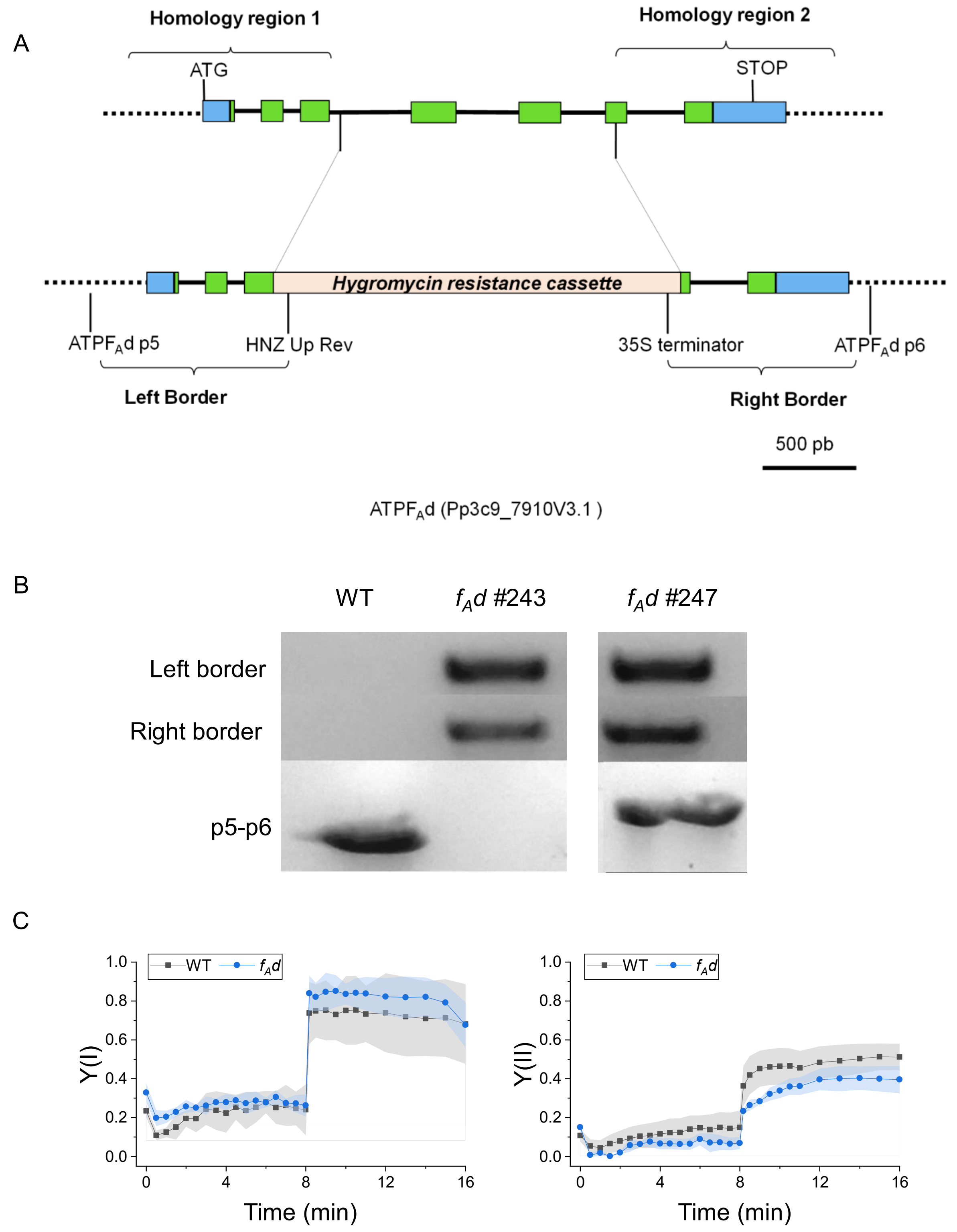


Figure S4 | **Generation of *P. patens* mutant lines lacking ATP synthase subunit F$_A$d and photosynthetic properties.** (A) Scheme showing the CDS, the regions of homology chosen to drive homologous recombination and insertion of the resistance cassette. (B) Example of PCR for verification of the homologous recombination event. PCR products called Left and Right Border are generated only if the resistance cassette is inserted in the expected genomic region. PCR product called p5-p6 is generated only if the cassette is not inserted (small fragment, WT) or inserted only once (larger fragment, #247). Line #243 lacks the p5-p6 product, meaning that the WT gene is missing, possibly through insertion in tandem of multiple repeats of the resistance cassette, a frequent phenomenon during gene targeting in *P. patens* [2]. (C) Yield of PSI and PSII in plants exposed to 330 µmol photons m$^{-2}$ s$^{-1}$ for 8 minutes, followed by 8 minutes in the dark.

**Table S1 | Identification of homologous sequences responsible for the formation and assembly of respiratory Complex V in eukaryotic organisms.** Protein and gene sequences from *Bos taurus*, *Saccharomyces cerevisiae* and *Chlamydomonas reinhardtii*, were obtained from ENTREZ at the National Center for Biotechnology Information (NCBI) server (http://www.ncbi.nlm.nih.gov/) while in *Arabidopsis thaliana* were obtained from the Arabidopsis Information Resource (http://arabidopsis.org/info/agi.jsp) or were identified using the PSI-BLAST tool available at the NCBI server. Names of proteins are based on published papers and reviews (see references). *P. patens* nuclear homologous sequences were identified using BLAST facilities with fungal, mammal, and plant protein and genic sequences against the *P. patens* genome (v3.3) (Lang et al., 2018; https://phytozome.jgi.doe.gov/pz/portal.html). Subunits encoded in the mitochondrial genome were identified from Terasawa et al., 2007 and GenBank accession numbers are given. Name for the subunits in plants are reported only when they differ from cattle or yeast [3, 4, 5, 6].

| *B. taurus* | *S. cerevisiae* | *A. thaliana* | *C. reinhardtii* | *P. patens* | Ref. |
|---|---|---|---|---|---|
| ***$F_1$ head*** | | | | | |
| ATP5A1/α | *α (ATP1)* | AtMg01190, At2g07698 | XP_001699641 | YP_539029 | 3,4,5 |
| ATP5B/β | *β (ATP2)* | At5g08670, At5g08680, At5g08690 | XP_001691632 | Pp3c5_26340, Pp3c5_26350, Pp3c6_4230 | 3,5 |
| ***Central stalk*** | | | | | |
| ATP5C1/γ | *γ (ATP3)* | At2g33040 | XP_001700627 | Pp3c18_8580, Pp3c19_11630, Pp3c19_5760, Pp3c21_11700 | 3,5 |
| ATP5D/δ | *δ (ATP16)* | At5g47030 | XP_001698736 | Pp3c4_31840, Pp3c13_11630, Pp3c26_13290 | 3,5 |
| ATP5E/ε | *ε (ATP15)* | At1g51650 | XP_001702609 | Pp3c1_10140, Pp3c1_12120, Pp3c2_24870 | 3,5 |
| ***Peripheral stalk*** | | | | | |
| ATP5F1/B | ATPB (ATP4) | AtMg00640 | – | YP_539002 | 3,4,5 |
| ATP5O/OSCP | ATP5 | At4g09650, At5g13450 | XP_001695985 | Pp3c11_16260, Pp3c21_15430, Pp3c23_11680, Pp3c24_12520 | 3,5 |
| ATP5H/D | ATPD (ATP7) | At3g52300 | – | Pp3c20_2830, Pp3c20_2850, Pp3c24_13710 | 3,5 |
| ATP5PF | ATPH (ATP14) | – | – | – | 5 |
| ***$F_O$ motor*** | | | | | |
| ATP6/A | ATPA (ATP6) | AtMg00410, AtMg01170 | XP_001689492 | YP_539022 | 3,4,5 |
| ATP5G3/C | ATPC (ATP9) | AtMg01080 | XP_001701531, XP_001701500 | YP_539041 | 3,4,5 |
| ATP5I/E | ATPE (ATP21) | At5g15320 | – | Pp3c1_20190, Pp3c2_20190 | 3,5 |
| ATP5L/G | ATPG (ATP20) | At2g19680, At4g26210, At4g29480 | – | Pp3c7_10, Pp3c17_14360 | 3,5 |
| ATP5J2/F | ATPF (ATP17) | At4g30010 | – | Pp3c5_15280, Pp3c5_15650, Pp3c25_5460, Pp3c25_5490 | 3,5 |
| – | APTI (ATP18) | – | – | – | 5 |
| – | ATPK (ATP19) | – | – | – | 5 |
| ATP8/A6L | ATP8 | AtMg00480 | – | YP_539003 | 3,4,5 |
| ***Plant specific*** | | | | | |
| – | – | At3g46430, At5g59613 (6Kda) | – | Pp3c12_2550 | 5 |
| – | – | At2g21870 (ATP7, $F_Ad$) | – | Pp3c9_7910 | 3, 5, 6 |
| ***Inhibitory Factors*** | | | | | |
| ATPI/IF1 | INH1, STF1 | At5g04750 (IFI-1), At2g27730 (IFI-2) | – | Pp3c1_22500 | 3,5 |

| | | | | | |
|---|---|---|---|---|---|
| **Assembly factors** | | | | | |
| ***$F_O$ subcomplex*** | | | | | |
| – | ATP10 | At1g08220 | – | – | 3 |
| ATP23 | ATP23 | At3g03420.1 | XP_001691633 | Pp3c20_19010, Pp3c24_12610 | 3 |
| – | ATP25 | – | – | | 3 |
| OXA1L | OXA1 | At5g62050 | XP_001693158 | Pp3c11_26830 | 3 |
| ***$F_1$ subcomplex*** | | | | | |
| ATPAF1 | ATP11 | At2g34050 | XP_001690396 | Pp3c14_19140 | 3 |
| ATPAF2 | ATP12 | At5g40660 | XP_001697254 | Pp3c2_19250 | 3 |
| – | FMC1 | – | – | | 3 |
